# Climate change in the Arctic: testing the poleward expansion of ticks and tick-borne diseases

**DOI:** 10.1101/2022.07.20.500759

**Authors:** Karen D. McCoy, Céline Toty, Marlène Dupraz, Jérémy Tornos, Amandine Gamble, Romain Garnier, Sébastien Descamps, Thierry Boulinier

## Abstract

Climate change is most strongly felt in the polar regions of the world, with significant impacts for the species that live in these extreme environments. The arrival of parasites and pathogens from more temperate areas may become a significant problem for these populations, but current observations of parasite presence often lack a historical reference of prior absence. Observations in the high Arctic of the seabird tick *Ixodes uriae* suggested that this species recently expanded its range poleward. As this tick can have a direct impact on the breeding success of its seabird hosts and is vector of many potential disease agents, including Lyme disease spirochaetes, its presence and origin are important elements for predicting its impact on polar seabird populations. Here, we use population genetic data and host serology to test the hypothesis that *Ixodes uriae* has recently expanded into the Svalbard archipelago. Both Black-legged kittiwakes (*Rissa tridactyla*) and Thick-billed murres (*Uria lomvia*) were captured in Kongsfjorden, Spitsbergen and sampled for ticks and blood. Collected ticks were genotyped using microsatellite markers and population genetic analyses were carried out using data from 14 additional seabird colonies distributed across the tick’s northern distribution. In contrast to predictions based on a recent expansion, the Spitsbergen population showed high genetic diversity and significant differentiation from the more southern populations, suggesting long-term population isolation. Host serology also demonstrated a high exposure rate to Lyme disease spirochaetes (Bbsl). Targeted PCR on tick DNA extracts and sequencing identified the presence of *Borrelia garinii* in a Spitsbergen tick, confirming seabird exposure and demonstrating the presence of Lyme disease bacteria in the high Arctic for the first time. Taken together, results contradict the notion that *Ixodes uriae* has recently expanded into the high Arctic region. Rather, this tick has likely been present for some time, maintaining relatively high population sizes and an endemic transmission cycle of Bbsl spirochaetes. Close future observations of population infestation/infection rates will now be necessary to relate epidemiological changes to ongoing climate modifications.

**Author summary:** The climate in the Arctic is rapidly changing, and with it, the flora and fauna that live there. These new environmental conditions can favor the establishment of invasive species, including novel parasites and pathogens. Here, we use population genetic data and host serology to examine whether recent observations of ticks infesting breeding seabirds in the high Arctic represent a poleward expansion of the parasite. Contrary to predictions, tick populations showed no evidence of a recent colonization of the region. Ticks have likely be present for a relatively long time, maintaining high local diversity despite harsh environmental conditions and vectoring infectious agents among breeding birds. Indeed, we demonstrate the presence of Lyme disease spirochaetes in the high Arctic for the first time, with bacterial DNA found in one of the sampled ticks and seabird serology demonstrating high exposure to this pathogen. This Lyme disease agent has therefore likely been established in the region and circulating at low frequency between seabirds and ticks for some time.

## Introduction

The climate is changing quickly and particularly so in the highly sensitive Arctic region where ice sheets are melting at an alarming rate and record high temperatures have become the norm (1,2). As a consequence of such changes, animal and plant distributions in these regions are shifting with potential cascading effects along trophic chains (3–5). Species that were previously mal-adapted to survive under polar conditions may now be able to invade, including novel parasites and pathogens (6,7). However, while there is no doubt about the potential impacts of climate change, precaution is required before declaring an emergence. Solid baseline data that support the absence of a parasite or pathogen is necessary in order to support claims of recent invasion. This data is often lacking, particularly for isolated regions and populations where few historical observations exist.

Recent studies suggest that ticks and Lyme disease are emerging in more northern areas due to climate change (8)(9), and becoming significant public health problems (e.g., (Hvidsten, Stordal et al. 2015). In polar regions, few tick species have been able to colonize, except those associated with colonial seabirds. These colonies represent particularly good environments for nidicolous ticks because host birds aggregate in high numbers to breed and return to the same nest site year after year in a very predictable manner (e.g., McCoy et al. 2016). However, at high latitudes, these ectoparasites also have to deal with harsh off-host conditions while waiting several months for their host to return for the next breeding season. Among tick species found in polar seabird colonies, the hard tick, *Ixodes uriae*, is the most wide-spread and abundant (10–12). *I. uriae* has evolved specific physiological and behavioural adaptations to deal with extreme conditions of temperature and humidity (13,14), but has been considered rare or absent in the high polar seabird colonies until relatively recently. Indeed, ticks were noted for the first time in penguin colonies of the Antarctic peninsula in the early 2000s, where it was suggested that their numbers were increasing (11). Population genetic studies indicated that these tick populations were not the result of recent colonisation or expansion events, but rather had existed in the area for a long period of time unnoticed by ornithologists (15). In the high Arctic, the presence of *I. uriae* was noted as early as 1999 in the Ossian Sarsfjellet colony of Spitsbergen (16) on a Black-legged kittiwake *(Rissa tridactyla*) chick, but at extremely low densities; only 1 larval tick was found on 67 captured birds, 61 kittiwakes and 6 Thick-billed murres (*Uria lomvia*). In 2007, Coulson and colleagues (17) observed ticks on Thick-billed murres in two colonies of the same area (prevalence of 20%), but did not find them in smaller murre colonies, nor on Black-legged kittiwakes. The authors suggested that *I. uriae* may have recently colonised the region, or had undergone a population expansion due to milder winter conditions or to weaken seabird immune responses caused by pollution.

From this time onward, specific efforts to record the presence of *I. uriae* in Svalbard have been made; Descamps (2013) found a clear relationship between the prevalence of ticks and the average temperature of the previous winter, supporting the notion that the establishment and spread of this tick is linked to environmental changes that favour tick survival. However, the question still remains as to whether the presence of this tick in the high Arctic represents a recent phenomenon, with its establishment and spread since the late 1990’s, or whether these populations have been present over longer periods of time, but were unobserved.

*Ixodes uriae* is a known vector of several pathogenic infectious agents (10), the most significant from a human perspective being the bacteria responsible for Lyme disease, bacteria of the complex *Borrelia burgdorferi* sl (also referred to as *Borreliella burgdorferi* sl). Indeed, Olsen and colleagues (18,19) showed early-on that *Borrelia garinii* spirochaetes could be found in *I. uriae* ticks sampled from temperate seabird colonies. Later serological studies demonstrated that colonial seabirds are frequently exposed to this infectious agent, particularly in North Atlantic colonies where the prevalence of seropositive birds averages almost 40% (20,21). Seropositive individuals and infected ticks have also been found within sub-Arctic colonies of northern Norway and Iceland (22–25). Genetic analyses of infected ticks demonstrated that the most frequently encountered Borrelia species in these seabird colonies is *B. garinii*, but other genospecies, such as *B. burgdorferi* ss and *B. afzelii* also occur (22,23). In the high Arctic, a PCR screening of ticks collected in Spitsbergen, Bjørnøya and Jan Mayen between 2008 and 2012, did not detect Borrelia spirochaetes (26), although sample sizes from Svalbard were extremely low (Spitsbergen = 9 ticks, Bjørnøya = 11 ticks). In the southern hemisphere, studies have demonstrated the presence of *Borrelia* in association with King penguins in the subantarctic islands (27), but have not detected Borrelia at higher latitudes (28).

Here, we test the hypothesis that *Ixodes uriae* has recently expanded into Spitsbergen, an Arctic island that has experienced severe temperature increases over the last three decades (29). We use a population genetic approach to compare the genetic diversity and structure of ticks sampled on Spitsbergen with those from more southern locations in the Northern hemisphere. If the arrival and spread of *I. uriae* represents a recent phenomenon, we expected to see low diversity within the Spitsbergen tick population relative to the other locations due to a founder effect. Using the genetic signatures from the southern colonies, we then determined the potential origin of the Spitsbergen population using tests of population structure and individual multi-locus assignments. If the establishment of *I. uriae* populations is recent, we also expected a low frequency of infectious agents in this population, as there should be a delay between the arrival of the vector and the arrival and spread of associated vector-borne agents (30–32). We examined this prediction in two ways: by analyzing the exposure of seabirds in Spitsbergen to *Borrelia* spirochaetes using serology and by the direct detection of bacterial DNA in the sampled ticks.

## Materiel and methods

### Study site, species and collections

Adults of two seabird species, the Thick-billed murre (*Uria lomvia*) and the Black-legged kittiwake (*Rissa tridactyla*), were captured during reproduction at the Ossian Sarsfjellet colony, Kongsfjorden, Spitsbergen, Norway (78°55’N 12°26’E) using a noose pole in 2012 and 2014. Both species have large colonies in the reserve, divided among several distinct cliffs where breeding takes place from May until August each year. Whereas kittiwakes breed in individual nests built on vertical cliff areas, murres aggregate and breed directly on cliff ledges. These birds are long-lived and highly faithful to both their breeding partner and breeding site among years (33). During capture, birds were searched for ticks and a blood sample (1 ml) was taken from the ulnar vein for serological analyses. This blood was centrifuged after sampling for approximately 10 mins at 5000 rpm and the plasma was extracted and stored at -20°C until immunological assays. All work was carried out in accordance with standard animal care protocols and approved by the Ethical Committee of the French Polar Institute (France) and the Norwegian Animal Research Authority.

*Ixodes uriae* (Family Ixodidae) is a colonial seabird specialist tick that occurs in polar regions of both hemispheres. Like other ixodid ticks, it has three developmental stages (larvae, nymph, and adult), but a relatively long life cycle (∼3 to 4 years) because of colder environmental conditions and limited host access (i.e., only during the relatively short breeding season). Larvae and nymphs of both sexes attach to the bird for a single blood meal that lasts from 3 to 8 days, after which they return to host nesting environment to moult and overwinter. As adults, only female ticks feed on the host (c.10 days) before laying a single clutch of several hundred eggs and dying (34,35). Mating typically occurs prior to the female blood meal in the nesting environment (36). In the present study, all captured birds were searched for ticks and any found were removed and conserved in 70% ethanol until analyses.

### Population genetic analyses

For ticks collected in the Ossian Sarsfjellet colony, DNA from each tick was extracted individually following the protocol of Kempf et al. (37). These samples were then genotyped at eight independent microsatellite markers developed specifically for this tick species (38). Microsatellite PCR amplification and allele size determination were performed as described in (37). Genotypes were visualized using an ABI PRISM 3130xl Genetic Analyser and allele sizes were assigned using GeneMapper v.4 (Applied Biosystems, Foster City, CA, USA).

For the other tick populations included in statistical analyses, we used the microsatellite data previously published by (39,40), along with two new population locations sampled in 2016 from Grimsey, Iceland and from Gull Island, Newfoundland, Canada (Figure 1). These locations were chosen to cover representative populations from the different possible source locations for ticks of the Ossian Sarsfjellet colony. In all locations, ticks were sampled from available host species, but ticks from different sympatric seabird species were treated as independent populations for the analysis based on results of previous work that demonstrated the presence of host-specific tick races within colonies (39,41–43). The three additional tick populations were genotyped as described above for the Ossian Sarsfjellet ticks, and control ticks from previously analysed populations were included to calibrate genotypes among datasets. The final data set used for analyzing the diversity and structure of the Svalbard population was therefore composed of 15 populations from six locations (Table 1). Although the included populations were sampled over a five-year period, we did not expect this to alter our biological inferences on population structure. The generation time of *I. uriae* is long (3 to 4 years) and therefore, at most, we have included data from two tick generations. Likewise, tick population sizes are typically large in most colonies and therefore resistant to rapid genetic drift. Indeed, here, we combined two different years for the Svalbard population, 2012 and 2014 (Table 1), but found no suggestion of between-year population structure (i.e., no heterozygote deficit suggested by the estimated F_IS_; see below and Table S1).

**Table 1:** Tick populations used in genetic analyses and diversity measures across loci. The colonies from the North Atlantic are indicated in orange, those from the North Pacific are in blue and those from Spitsbergen (Svalbard) are in green. Raw genotype data are available in Table S3.

| Colony location | Year sampled | Seabird species* | No. ticks (No. hosts) | Avg allelic richness (±SE)** | Avg gene diversity (±SE) |
| --- | --- | --- | --- | --- | --- |
| Hornøya (HN), Norway | 2010 | TBMU | 35 (21) | 3.88 (±2.15) | 0.52 (±0.26) |
| | 2009 | COMU | 33 (off†) | 4.66 (±2.44) | 0.59 (±0.26) |
| | 2009 | BLKT | 35 (-#) | 4.85 (±1.76) | 0.66 (±0.21) |
| | 2009 | ATPU | 35 (off†) | 4.90 (±2.85) | 0.57 (±0.32) |
| Grimsey (GM), Iceland | 2016 | COMU | 30 (off†) | 4.81 (±1.82) | 0.65 (±0.22) |
| | 2016 | ATPU | 28 (off†) | 4.31 (±2.71) | 0.50 (±0.37) |
| Gull Island (GU), Canada | 2016 | ATPU | 19 (off†) | 4.92 (±3.14) | 0.61 (±0.32) |
| St. George (SG), Alaska | 2009 | TBMU | 20 (20) | 5.15 (±2.82) | 0.63 (±0.29) |
| | 2009 | COMU | 31 (20) | 5.32 (±2.94) | 0.64 (±0.24) |
| | 2009 | BLKT | 17 (14) | 4.42 (±2.12) | 0.58 (±0.27) |
| Bogoslof (BO), Alaska | 2009 | TBMU | 28 (13) | 5.39 (±2.81) | 0.63 (±0.28) |
| | 2009 | COMU | 27 (12) | 5.42 ($\pm 2.72$) | 0.65 ($\pm 0.27$) |
| | 2009 | BLKT | 27 (11) | 5.27 ($\pm 2.91$) | 0.66 ($\pm 0.20$) |
| | 2009 | TUPU | 16 (>2††) | 4.74 ($\pm 2.08$) | 0.67 ($\pm 0.19$) |
| Ossian Sarsfjellet (OS),<br>Svalbard | 2012/14 | TBMU | 37 (37) | 4.06 ($\pm 1.70$) | 0.57 ($\pm 0.20$) |

\*TBMU=Thick-billed murre; COMU = Common murre; BLKT = Black-legged kittiwake; ATPF= Atlantic puffin
(Atlantic Ocean only); TUPU=Tufted puffin (Pacific Ocean only)
\*\*based on 8 randomly sampled tick individuals in each population
† off = all ticks off-host from around nesting; # gorged ticks were collected from over 50 birds, but host identity was lost post-moult and therefore the true number of individual host birds is not available; ††one tick collected on-host, all others from the off-host environment

**Figure 1.**
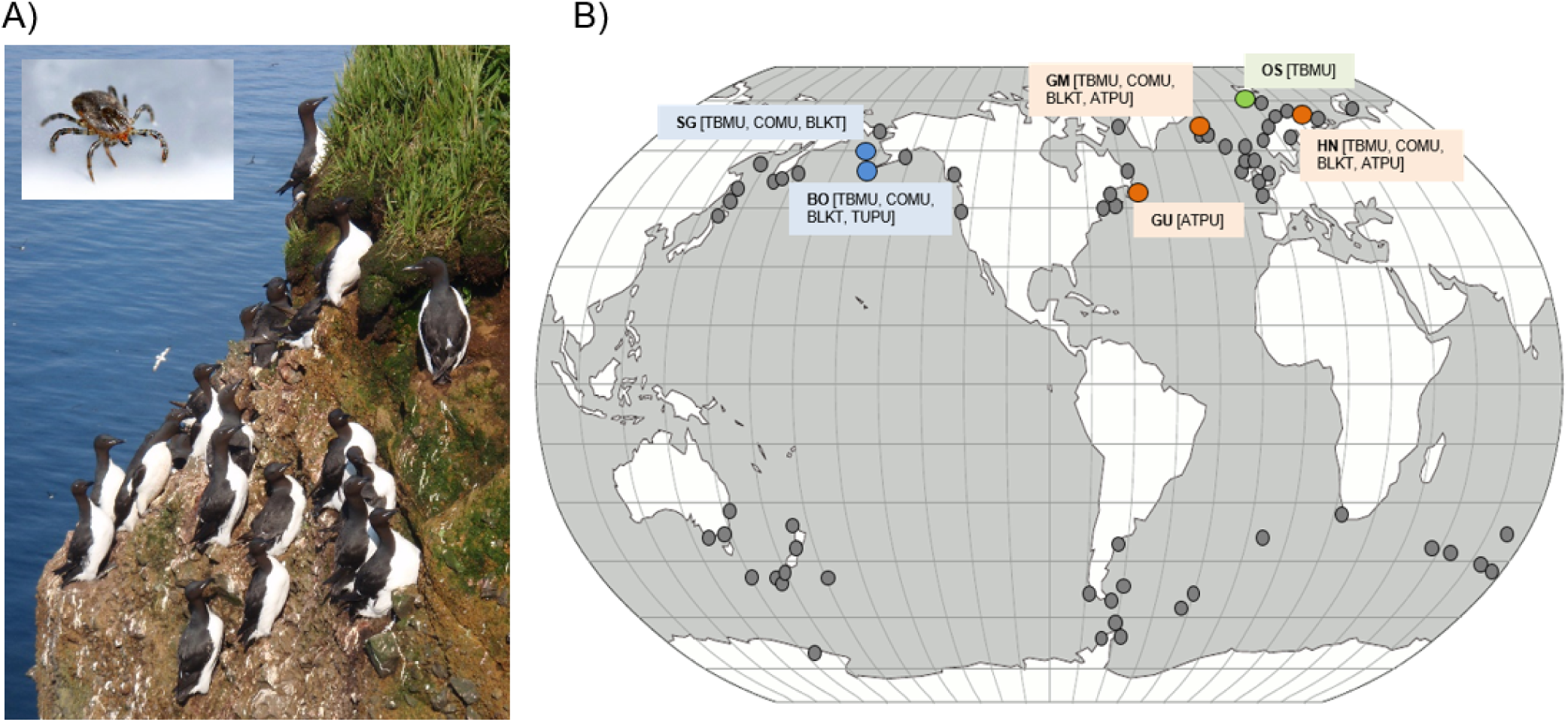
A) *Ixodes uriae* (inset) exploits seabird species such as Thick-billed murres *Uria lomvia* across the polar regions of both hemispheres. B) The known worldwide distribution of *Ixodes uriae*, with locations where the tick has been directly observed indicated by grey points (after (10,12). The central study population in Ossian Sarsfjellet, Svalbard is indicated by a green point and the five additional locations used for comparative population genetic analyses are indicated in blue (North Pacific locations) and orange (North Atlantic locations) with seabird host species sampled for ticks indicated for each location. Acronyms for locations and seabird species are given in Table 1. Photo credits: P. Landemann (*I. uriae*); K.D. McCoy (*U. lomvia*).

Genetic diversity in each colony was estimated by calculating allelic richness (AR) using a rarefaction method, and Nei’s unbiased gene diversity (H_S_). Within-population departures from Hardy–Weinberg (HW) proportions were investigated by estimating the inbreeding coefficient (F_IS_). The significance of this estimator was assessed by randomizing alleles among individuals within samples (2400 permutations). These basic analyses were all carried out using FSTAT (44).

To examine population isolation, pairwise F_ST_ estimates were calculated from microsatellite data according to Weir and Cockerham (45). Significance was assessed by permuting multi-locus genotypes among populations using FSTAT (44), with 2100 permutations and a significance level of 5%, corrected using the standard Bonferroni method for multiple tests.

Population genetic structure was also inferred using a Bayesian clustering approach implemented in the program STRUCTURE 2.3.4 (46). The program was run with five independent runs, using the admixture model with correlated allelic frequencies and the LOCPRIOR model developed in Hubisz et al. (47) to take into account the sampling location of each individual. All simulations used 100 000 iterations in the burn-in phase and 100 000 generations in the data collection phase. Selection of the number of distinct clusters (k) was based on Evanno’s DeltaK (48) using the Harvester software (49). STRUCTURE analyses were conducted on two partitions of the data: (i) the entire data set to identify those populations that were most related to the tick population on Spitsbergen (n = 418 ticks from 15 populations, testing k = 1 to 15), and (ii) the three most closely related populations to examine finer scale structure (n = 135 ticks from four populations, testing k = 1–5).

### Immunological assays

Plasma from 19 Black-legged kittiwakes and 16 Thick-billed murres sampled in 2014 from Ossian Sarsfjellet were screened for antibodies directed against three subspecies of *Borrelia burgdorferi* sensu lato (Bbsl; *B. garinii, B. afzelii* and *B. burgdorferi* ss) using a commercially available enzyme-linked immunosorbent assay (ELISA) kit (Borrelia + VIsE IgG ELISA, IBL International, Hamburg, Germany). This kit was slightly modified for use on seabirds (as per Staszewski et al., 2008). Briefly, we replaced the secondary (anti-IgG) antibody of the kit by an anti-chicken IgY antibody conjugated with peroxidase (Sigma A-9046, Sigma-Aldrich) diluted to 1:750, and then followed the kit manufacturer instructions. Each sample was run in duplicate and anti-Bbsl antibody levels were quantified as the mean optical density (OD) of the resulting solution read at 450nm in a spectrophotometer (Victor 1420, Perkin Elmer). We used the method described in Garnier et al. (50) to determine negative and positive thresholds. As anti-Bbsl antibodies are known to persist for several years in seabirds (51), a seropositive status reflects an exposure event that occurred at an unknown time prior to sampling.

### Borrelia *detection in ticks*

We applied a highly sensitive target-specific qPCR procedure to detect the presence of *Borrelia* infection in ticks collected from the Ossian Sarsfjellet population following Gómez-Díaz et al. (52). Any positive samples were reamplified using the nested PCR procedure described in Duneau et al. (23) and were sent for direct sequencing (Eurofins). Sequences were cleaned and aligned using the ClustalW algorithm implemented in MEGA v7 (53) and an blastn search was carried out to determine species identity.

## Results

### Tick genetic diversity

Genetic diversity was relatively high in all populations, varying from 3.88 (±2.15) to 5.42 (±2.72) in allelic richness, and from 0.50 (±0.37) to 0.67 (±0.19) in gene diversity; the Svalbard tick population fell within the middle ranges of both estimates (Table 1). Tests of HW indicated that several tick populations did not conform to expectations (Table S1). In North Pacific populations, this was largely due to heterozygote deficits at locus T44. Previous studies have suggested that this locus may be linked to a gene under host-associated selection (39,43). When this locus was removed, all populations except ATPU-GU were shown to respect HW proportions. In the ATPU-GU population, the overall deficit was due to locus T39, and is likely linked to a combination of genotyping errors and low sample size (see Table S1). However, to account for potential non-independence in the data, tests for population differentiation did not assume HW equilibrium. In addition, STRUCTURE analyses were run with and without T44. As results were similar in both cases, only results that include all eight loci are presented below.

### Tick population genetic structure

Pairwise tests of population structure showed that the Spitsbergen population differs strongly and significantly from all other tick populations, with F_ST_ estimates varying from 0.0853 to 0.3843 (Table 2). These distances were lower overall than those between North Atlantic and North Pacific populations, but higher than those within the North Pacific and of similar magnitude as those within the North Atlantic (Table 2).

**Table 2.**
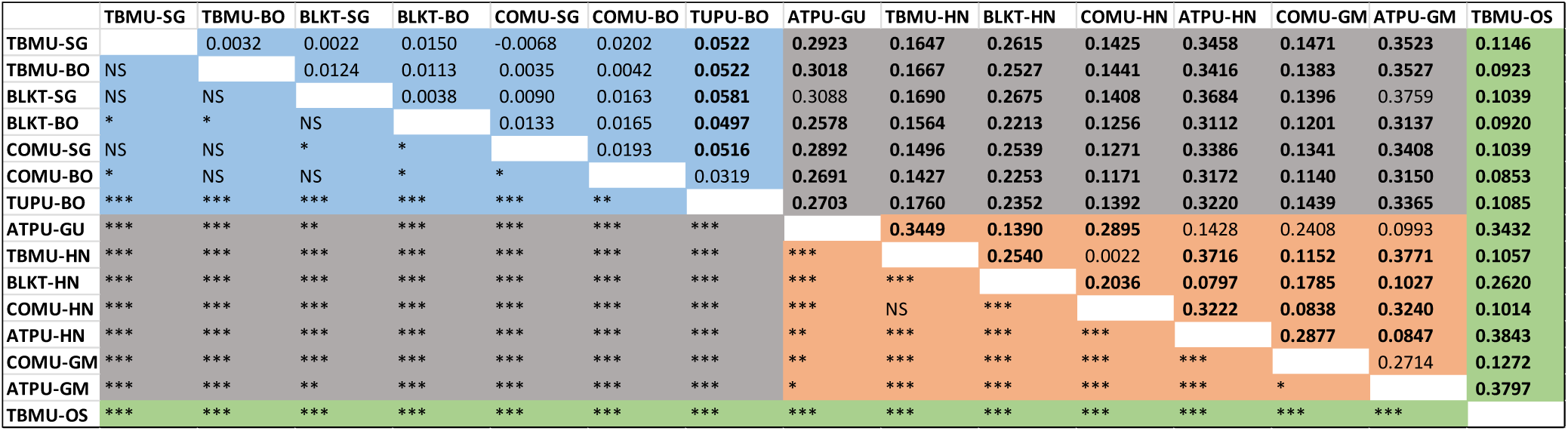
Pairwise population genetic structure among tick populations (n= 15 populations). Comparisons within and among ocean basins are highlighted in different colours (orange= North Atlantic seabird colonies, blue=North Pacific colonies, grey= North Atlantic versus North Pacific) and comparisons with the Spitsbergen population are shown in green. Estimates of F_ST_ are shown above the diagonal, with the associated significance levels below the diagonal. F_ST_ values in bold are those considered significant at the adjusted nominal level of 5% after accounting for multiple comparisons (P< 0.0005). The significance levels below the diagonal are as follows: *** P<0.0005, ** P<0.001, * P<0.05, NS P>0.05. Populations are identified by acronyms that indicate the seabird host species and the geographic location as outlined in Table 1.

Results from the STRUCTURE analysis mirrored tests of pairwise population structure. When all 15 populations were included in the analysis, two major clusters were identified according to the deltaK (see Figure S1), one in the Pacific, which included Atlantic murre ticks, and the other with the remaining Atlantic tick populations. The selection of k=2 is not surprising given the strong differentiation between ticks from different ocean basins (39,40) (Table 2). However, small peaks in deltaK values were also seen at k=3 and k=5 (Figure S1). When these clusters are examined in more detail, isolation of the Spitsbergen population becomes more apparent; when k=3, the Spitsbergen population groups with other murre ticks from the North Atlantic and when k=5, isolation of the Spitsbergen population becomes evident (Figure 2a). When STRUCTURE was ran a second time, but including only the three most closely related murre tick populations from the North Atlantic, the optimal number of clusters was 3 (see Figure S1). In this case, the Spitsbergen tick population was clearly isolated from the murre tick populations on Hornøya (Norway) and Grimsey (Iceland); no misassignments that might indicate recent gene flow are evident (Figure 2b).

**Figure 2.**
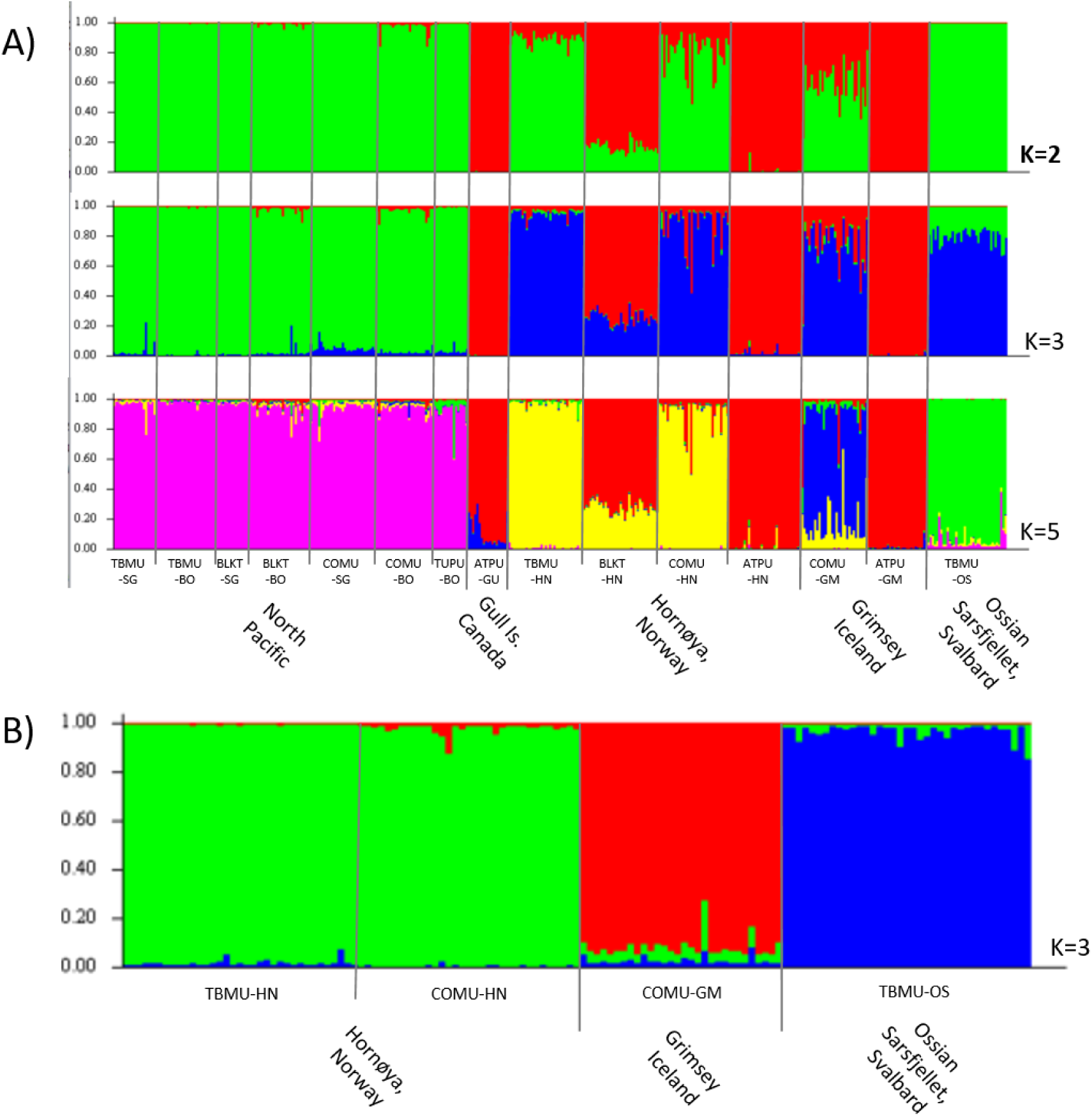
A) Multi-locus assignments of ticks to population groups when considering the 15 analysed tick populations and an increasing number of defined clusters (k=2, 3 and 5) according to STRUCTURE. Each individual bar represents the probability that a tick belongs to a cluster of a given color. The Ossian Sarsfjellet tick population groups with other ticks exploiting murres in the North Atlantic when K=2 (optimal number of clusters as defined by the deltaK: see text), but becomes more isolated as the number of defined clusters increases. **B)** Results when only the three more closely related tick populations to the Ossian Sarsfjellet colony are considered in the analysis. In this case, the optimal number of defined clusters was 3, with no indication of any recent tick migrations into the Ossian Sarsfjellet colony (ie, misassignments).

### Immunological assays

The plasma of 19 Black-legged kittiwakes and 16 Thick-billed murres were screened for Bbsl antibodies. The distribution of mean OD values based on a reference sample was bimodal for both species (Figure S2), with high OD values corresponding to seropositive plasma samples and low OD values to seronegative samples. Based on these defined threshold values for positive and negative tests (Table S2), 63% of sampled Black-legged kittiwakes were seropositive (12/19), with one ambiguous status, whereas all Thick-billed murres were seropositive (Figure S2).

### Borrelia detection

Of the 37 tick DNA extracts tested by qPCR, one was found positive for Bbsl infection. After reamplification and Sanger sequencing, we obtained a 379bp fragment that was used in the blastn search (Genbank accession number ON979421). The top 100 sequences identified in the blast search all corresponded to *B. garinii* (or a non-identified *Borrelia*) with more than 98.42% identity.

## Discussion

Climate change is rapidly altering polar environments, opening the door to invasive species, including novel parasites and pathogens (54). It was hypothesized that *Ixodes uriae* has recently expanded in the Svalbard archipelago due to these changing climatic conditions (17). Here, we examined the hypothesis of a recent expansion using a population genetic approach and explore the possible concomitant presence of Lyme disease spirochaetes at very high latitudes. The first documented observation of *I. uriae* in Svalbard to our knowledge is from 1999, when a single larval tick was found on a nestling in Kongfjorden (McCoy 2001). If this observation was due to the recent establishment of the tick, we would expect to see a strong founder effect signal a decade or so later (i.e., after roughly three tick generations), when sampling for our study took place. This was not the case. Here, we found that genetic diversity within the Spitsbergen tick population was equal to or higher than that of more southern populations. We also found that although this population was most closely related to that of the same seabird host species from populations of Norway and Iceland (i.e. murre hosts), it was still significantly isolated from these other populations, suggesting limited ongoing gene flow.

The presence of *I. uriae* in a seabird colony depends on three factors: the availability of a suitable host for the bloodmeal, an appropriate off-host environment for moulting and overwintering, and a means of transportation to colonise the location. Although considered a seabird generalist, *I. uriae* has been shown to repeatedly form host-associated races throughout its distribution (37,40,41,43). Therefore, the exploitation of one seabird species within a colony can be independent of that of other sympatric bird species, and depends on potential spillover dynamics (40). We can see in our results that only ticks collected from the closely related Thick-billed and Common murres showed no structure within colonies (Table 2), but some introgression among local host-associated tick populations is evident, demonstrating that host switching can occur (Figure 2). Thus, as several suitable avian host species breed in large numbers within the Svalbard archipelago, it is unlikely that the host *per se* represents a barrier to tick colonisation in this part of the world. Indeed, our results show that both Thick-billed murres and Black-legged kittiwakes in the Ossian Sarsfjellet colony are exposed to enough ticks and for a sufficient amount of time to be infected with Borrelia spirochaetes (see below). If the requirement for suitable hosts is met, the other two factors required for tick establishment are less evident.

In terms of the off-host environment, it was previously suspected that the cold, dry winters of the high latitudes were likely unsuitable for overwintering ticks (16,17). A recent experimental study of tick physiology demonstrated that temperature and humidity interact to determine *I. uriae* survival and that different life stages and physiological states are more or less susceptible to low temperatures (13). A regular survey of murres in the Kongsfjorden area started in 2005, with between 10 and 100 adult birds captured per year. *I. uriae* was first noted in 2007 (17) and targeted screening for ticks on captured birds started after this point. Prevalence in captured adult birds varied between 0 and 35% over the following six years of study. A correlation between the average temperature the previous winter and tick prevalence the following summer was highly significant, with winter temperature explaining almost 90% of the variance in tick prevalence over time. This correlation translated into a 5% increase in prevalence for every 1°C increase in winter temperature. However, no overall increasing trends in tick prevalence or in temperature were evident across the six years (55). It is therefore probable that tick population sizes increase with increasingly mild winters, and that the high Arctic is becoming more suitable for this tick. Indeed, surveys in a murre colony of Isfjorden (Diabasodden) recorded the presence of *I. uriae* for the first time in 2020 (S. Descamps, unpublished data). However, long-term trends in tick abundance remain difficult to predict as extreme events may have a major impact on tick survival (13). What is surprising and important in our results is that tick genetic diversity within the Ossian Sarsfjellet colony was similar to that found at lower latitudes. This means that, even if fluctuating winter conditions alter tick survival probability, enough ticks survive each year to maintain high local diversity. A prolonged life cycle that includes years of dormancy, when ticks to not attempt to feed and remain protected within the cliff environment, may help explain these patterns (56); changes in climatic conditions in certain years may then favour tick reactivation.

Finally, in order to colonize a new location, ticks must arrive at the site. As seen in the overall population genetic analyses presented here, structure tends to be high among populations in different locations, but depends on the tick host race. Indeed, dispersal in *I. uriae* is considered a relatively rare event because it depends on the movement of infested birds between breeding locations at a time when few birds move (57). It has therefore been suggested to occur during prospecting, when failed breeders or juvenile birds visit new colonies in late reproduction to choose a future breeding site (57,58). The probability of dispersing ticks during such visits depends on the behaviour of the different seabird species within the colony (42). For example, geographic structure among puffin tick populations within the North Atlantic is very weak compared to that in ticks associated with other species (McCoy et al., 2001, 2005; this study). Puffins nest in burrows or rock piles on cliff slopes and spend a good proportion of their time interacting outside the nest site (59). Visiting birds may land on a puffin slope without aggression and can therefore pick up or leave ticks relatively easily. This is less the case for highly territorial species such as murres, where visiting birds must remain at the outskirts of breeding ledges or suffer attacks from nesting individuals. This territorial behaviour should reduce the probability of tick dispersal among sites. In addition, murre populations in Svalbard have been declining since the 1990s (60) which may reduce prospecting in these colonies, and therefore the probability of birds carrying ticks into the area. In agreement with these observations, our genetic results suggest that ticks of the Ossian Sarsfjellet colony have been isolated for some time, with little current gene flow from other colonies.

Additional results from our study support the long-term presence of *I. uriae* in Svalbard and, in particular, the high prevalence of exposure to a tick-transmitted infectious agent, Lyme disease bacteria. Indeed, if the arrival of this tick was a relatively recent event, we would expect a lag in the arrival and spread of tick-borne pathogens (31). *I. uriae* is a known vector of Lyme disease spirochaetes, and populations frequently show high prevalence [e.g., 11% in kittiwake ticks (22)]. Correspondingly high exposure rates in diverse seabird species have also been observed [40% on average in North Atlantic colonies; (21); up to 18% in North Pacific colonies (20)]. Here, we found that all Thick-billed murres of Ossian Sarsfjellet were seropositive, demonstrating that birds are exposed to enough ticks to eventually become infected and mount a humoral immune response. Exposure rates in kittiwakes of this same colony were lower (63%), showing that kittiwakes are also exposed to ticks within this colony. This data, combined with the high diversity and isolation of the tick vectors, suggests that Borrelia is maintained in an enzootic cycle in the Ossian Sarsfjellet seabird community.

To verify the presence of Lyme disease spirochetes in ticks from Spitsbergen, we carried out targeted searches for Borrelia in tick DNA extracts and demonstrated the presence of a *B. garinii* isolate. Despite previous attempts (26), this is the first demonstration of Lyme disease spirochetes in the high Arctic. Only one tick of 37 tested were found to be positive using our qPCR protocol (ie, prevalence <3%). This prevalence, combined with relatively low observed infestation levels, renders high seroprevalence in the birds difficult to explain. The apparent contrast may arise from several sources. First, ticks were extracted and tested after several years of storage in ethanol, potentially reducing the ability to detect harboured pathogens. In addition, a previous study showed that Borrelia spp. detection probability was lower in murre ticks compared to ticks from other host races due to lower infection intensities (52). Estimated prevalence of Borrelia in ticks may therefore be underestimated. Second, seropositivity reflects an exposure event at an unknown time in the past because antibodies are maintained over several years (51). Among seabird species, Borrelia seroprevalence of murres has been found to be higher than in other sympatric species, suggesting that these birds may react more strongly to infection and/or maintain specific antibodies for longer periods of time (21,61). Indeed, here we found that optic density values (a proxy of antibody levels) in seropositive individuals covered a range values, suggesting a mixture of recent and past exposure to Borrelia (Figure S2). Finally, there can be high heterogeneity in the presence of infected ticks at the colony scale (22); in our study, ticks were mostly sampled in 2012, whereas serology was carried out on birds captured in 2014. Samples may therefore have come from slightly different parts of the breeding colony.

Polar environments are changing quickly, with important consequences for the species that live there. As the poles warm, new species can invade, potentially adding stress to populations that are already dealing with changes in food resources, breeding habitats and increasing pollution. The invasion of novel parasites and pathogens is particularly important to survey in this respect. We show here that contrary to previous suggestions, *I. uriae*, and at least one of its associated pathogens, do not seem to be new additions to high Arctic parasite fauna. However, if infestation levels have been relatively benign in the past, climate change may alter this. Increasing temperatures may enable higher tick survival, and thus result in higher parasite pressure on breeding birds and their young and higher exposure rates to tick-borne pathogens. A warmer Arctic may advance the timing of breeding, and thus may alter the nature of tick exploitation in terms of the avian species or the life stages (adult or chick) being exploited. Additional parasites and pathogens may also be introduced and act in synergy with ticks. Extreme climate events may limit some of these effects by periodically lowering parasite survival in the environment. Only continued monitoring of these polar populations will enable us to evaluate these different possibilities and to better relate on-going epidemiological patterns to climate change.

## Acknowledgements

We thank the following people for their assistance with sampling seabirds and ticks: Delphin Ruche, Saga Svavarsdottir, and Pierre-Axel Monternier (Spitsbergen); Thorkell Lindberg Thórarinsson and Yann Kolbeinsson from the North East Iceland Nature Center (Iceland); Gregory Robertson and David Fifield from Environment and Climate Change Canada (Newfoundland, Canada). We also acknowledge help from Roger Colominas-Ciuró, and Quentin Schull for serological analyses, and from Muriel Dietrich and Valérie Noel for microsatellite genotyping. The capture of seabirds and removal of ectoparasites in Ossian Sarsfjellet was approved by the Governor of Svalbard (program number 361 and permit 2014/731).

Financial support for sampling was provided by the French Polar Institute (IPEV, project no. 333) and the Agence National de la Recherche (ANR-13-BSV7-0018-01). M.D. and A.G. were supported by fellowships from the French Ministry for Education and Research.

